# Deciphering the developmental program in the ascidian *Ciona intestinalis* just prior to gastrulation

**DOI:** 10.1101/137166

**Authors:** Yoshito Harada

## Abstract

The embryo of the ascidian *Ciona intestinalis* displays striking anatomical simplicity, with an invariant cleavage pattern during development. It has a monoploid genome like the model organisms *Drosophila melanogaster* and *Caenorhabditis elegans*, whereas vertebrates generally exhibit tetraploidy. In this study, I took advantage of these characteristics to investigate the development of the organism as one reverse-engineers an industrial product. First, the spatial expression of 211 genes was digitalized. Some genes showed variable expression patterns, which might reflect multiple snapshots of a single temporally dynamic expression at different times. Several developmental territories of the embryo were considered to be very similar to each other; however, this digitalization of gene expression patterns showed that differences occurred between individual blastomeres even within a single developmental territory. Furthermore, *Ciona Brachyury* (*Ci-Bra*) was expressed in those blastomeres in which both *Fox (forkhead-box) A-a* and *Zic (zinc-finger) L* were expressed, these proteins being upstream regulators of *Ci-Bra.* The approach described enables the developmental program to be studied *in silico*.

**SUMMARY STATEMENT:** A detailed expansion of our knowledge on an animal developmental program by using already published gene expression data with the aid of computers

## INTRODUCTION

Studies on the developmental programs of model organisms such as the fruit fly *Drosophila melanogaster* and the nematode *Caenorhabditis elegans* have furthered our knowledge greatly (Lawrence, 1992; WormBook: http://www.wormbook.org). In these animal models, saturation mutagenesis studies have been undertaken to screen for genes whose mutation causes abnormal morphological phenotypes. Subsequently, the hierarchical relationships among these genes have been determined and the underlying genetic sequences isolated. Consequently, hypothetical diagrams of the developmental programs of these organisms have been successfully constructed.

However, similar approaches in vertebrate animal models have not achieved the same success. Genes in vertebrates have undergone two rounds of duplication at the chromosome level; consequently, every gene exhibits fourfold redundancy with “ohnolog” paralogous genes (Ohno, 1970). The vertebrate gene regulatory network (GRN) has evolved from this initial state of fourfold redundancy to the current state, in which redundant subcircuits have been pruned off in each evolutionary lineage. For example, there are four *izumo* ohnologs (*izumo-1, 2, 3 and 4*) in the mouse genome, only one of which, the single *izumo-1* gene, results in infertility when knocked out (Inoue et al., 2005; Grayson and Civetta, 2012). Two scenarios might explain this situation (Fig. 1). The first is that an ancestral gene for the four *izumo* ohnologs, *proto-izumo*, played an essential role in fertilization and that the other three ohnologs (*izumo-2, 3 and 4*) lost the ability to complement *izumo-1* fully in an ancestor of mice. The second is that only the ancestor of the *izumo-1* gene was recruited for this fertilization function, and not the ancestors of the other ohnologs. There is no definitive answer as to which scenario is correct at present.

**Fig. 1.**
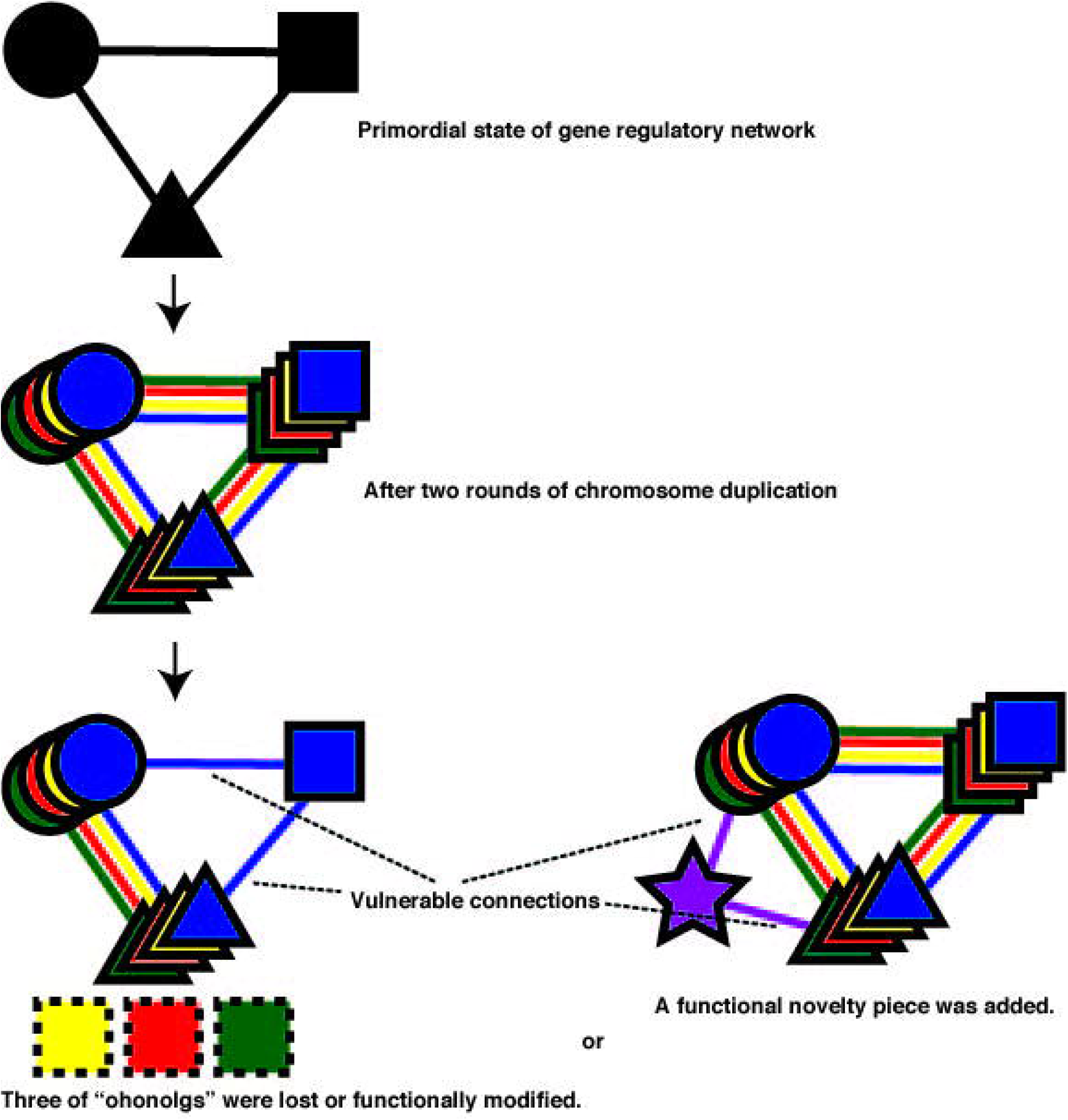
Two rounds of duplication of entire chromosomes in vertebrates. Two scenarios explain why in many genes a single mutation can cause phenotypic abnormalities, whereas every gene possesses quadruple redundancy in principle. (Left) The function of the original primordial gene is inherited by only one of the four ohnolog paralogues that arose during the two rounds of chromosome duplication. (Right) An innovative function is assigned to another accidentally appointed gene.

In vertebrates, a gene knockout phenotype might not necessarily reveal the ancestral biological function of the gene. A developmental regulatory gene in which a single hypomorphism causes an apparent abnormality represents a fragile point within the overall GRN (Fig. 1). Living organisms tend to exclude wastefulness from their genomes. Consequently, a paradoxical phenomenon might emerge: genes that are more critical for development might show fewer abnormalities than less important genes when they are knocked out because of functional redundancy with their backup copies.

The aim of the study described herein was to understand the encrypted developmental program of early embryonic development in the ascidian *Ciona intestinalis*. The ascidian tadpole larva represents the basic body plan of all chordates in a relatively small number of cells and tissue types. Extensive whole-mount *in situ* hybridization and experimental data on its molecular developmental mechanisms have been amassed. Each cell of the embryo possesses a restricted developmental fate. For example, mesenchyme cells are derived exclusively from only two blastomeres of the 110-cell embryo. The ascidian larva does not construct a typical archenteron such as that seen in the sea urchin or frog, instead the ascidian archenteron only forms a shallow dent. The most remarkable advantage of the ascidian tadpole larva as a model organism to elucidate the chordate developmental program is that it has a monoploid genome and therefore is free from the gene-redundancy problem as stated above.

When GRNs are compared among vertebrates, differences can be observed. For example, in the case of endoderm formation in *Xenopus laevis* (frog) and zebrafish, the GRN is not completely conserved between the two organisms (Shivdasani, 2002). Instead, they share only some key genes, which may be considered vertebrate endoderm master genes (genes that determine cells to differentiate into endoderm). One such gene is *Sox17*, which belongs to a family of transcription factors that contain the high mobility group box found in the sex-determining region Y gene. It is possible that the differences between the frog and zebrafish are caused by a simple technical artifact. However, it is also probable that in these species, redundant branches have been pruned independently from the quadruple prototype GRN. In the latter scenario, the aim of comparative studies of the frog and zebrafish GRNs is not to ratify the common elements of both networks but to discover new master genes such as *Sox17* to increase the number of genes that can be used for evolutionary developmental biology (evo-devo) studies.

The definition of GRN seems to have changed since its first introduction (Davidson, 2006), from the core network described in the first evo-devo studies (e.g. Halder et al., 1995) to incorporate the concept of subcircuits in 2007 (Hinman and Davidson, 2007; Davidson, 2010). The core network is a gene network that coordinates the differentiation of a relatively large embryonic territory (e.g. mesoderm) that develops into multiple cell types (e.g. muscle, mesenchyme and notochord). A subcircuit is an individual part of the whole GRN (e.g. the GRN that underlies muscle formation). The developmental area that is controlled by an individual subcircuit corresponds roughly to the area that is controlled by one of the aforesaid master genes.

The aim of the study described herein was to understand the encrypted developmental program of early embryonic development in the ascidian *C. intestinalis*. To elucidate the early embryonic developmental program of *C. intestinalis*, I developed two small computer programs (Encoder and Decoder) that digitalized microscope images of whole-mount *in situ* hybridization analyses (Fig. 2; Harada, 2016). For each gene analysed, a score of 1 or 0 was assigned to each blastomere depending on whether gene expression was ON or OFF. These scores then formed a character string for each embryo examined (the Encoder tool). The Decoder tool is based on the fact that multiple blastomere images with a transparent background can be stacked to provide a representation of expression throughout the vegetal half of an embryo. Both the Encoder and Decoder programs are freely downloadable from the figshare repository (https://figshare.com).

**Fig. 2.**
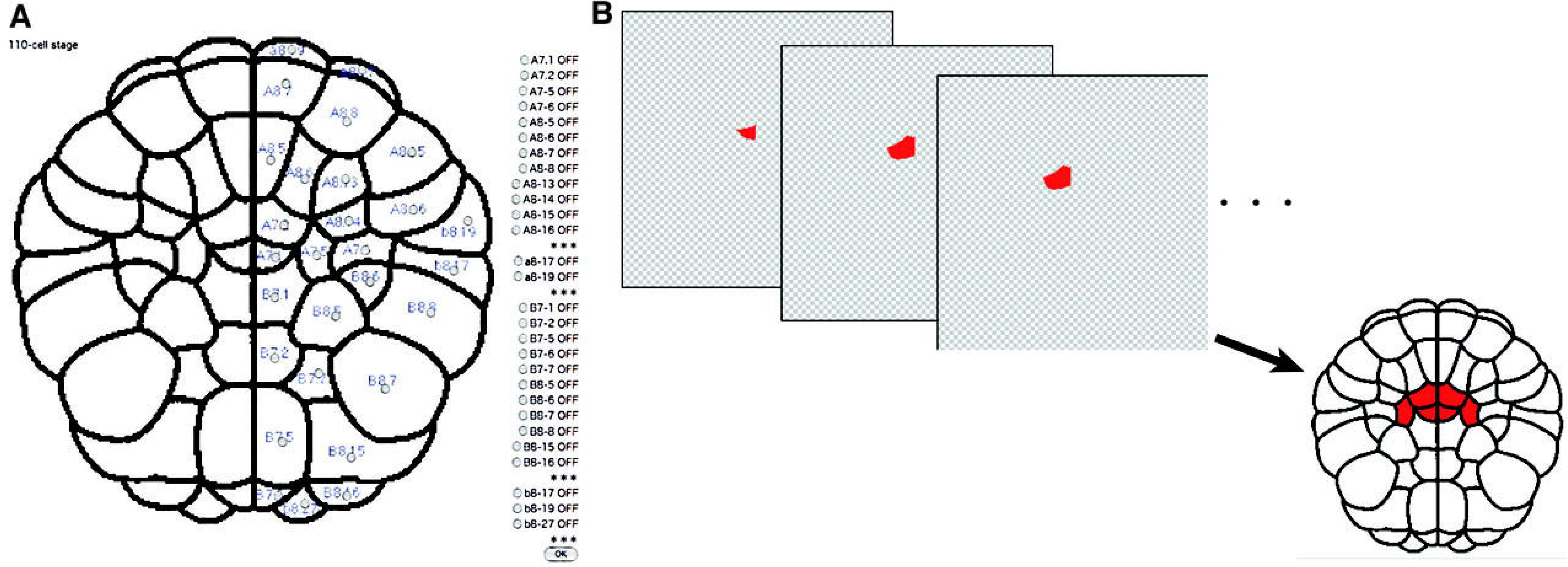
Encoder and Decoder programs. (A) Encoder program. (B) Decoder program. See the text in the Materials and Methods section for an explanation of these programs.

In various molecular studies of developmental biology to date, developmental territories have been shown to adopt an abnormal phenotype when the expression of a certain gene is increased or decreased. For example, ectopic *MyoD* activity causes extra muscles to develop in blastomeres that normally develop into ectoderm (Hopwood and Gurdon, 1990). However, even the four primordial muscle blastomeres (B8.7, B8.8, B8.15 and B8.16) of *C. intestinalis*, which are considered to have almost uniform developmental properties, it was proven that they exhibit some differences when gene expression is examined carefully (see below) although the true meaning of this difference remains unknown.

It is not feasible to perform such detailed studies in every developmental biology paper because they involve the reporting of the expression patterns of some tens of whole-mount *in situ* hybridization probes one by one. Instead, I propose that a data repository should be used to accumulate high-quality data for spatial patterns of gene expression at the cellular level during normal development. This would obviate the need for researchers to perform experiments *in vivo* or *in vitro*; instead, they could examine their hypotheses virtually (*in silico*) using the repository of published data. This paper illustrates the utility of this approach in the chordate *C. intestinalis*.

## RESULTS

### Characterization of gene expression patterns in Ciona intestinalis embryos using the Encoder and Decoder tools

I aimed to decipher the developmental programs encrypted in the *C. intestinalis* genome. Using the Encoder tool, I digitalized the spatial expression patterns of 211 *C. intestinalis* genes from the Ascidian Network for *in situ* Expression and Embryological Data (ANISEED) development browser (Tassy et al., 2010; Brozovic et al., 2016; http://www.aniseed.cnrs.fr) into character strings (Figs. 2–4). It is known that three maternal factors, β-catenin, Zic-r.a (macho-1) and GATA.a. are required to establish the zygotic expression patterns of *C. intestinalis* genes that give rise to the vegetal hemisphere, posterior vegetal hemisphere and animal hemisphere, respectively (Satoh, 2013; Ikeda and Satou, 2017). The animal hemisphere comprises the descendants of the a- and b-line blastomeres, whereas the anterior vegetal hemisphere comprises the descendants of the A-line and the posterior vegetal hemisphere comprises the descendants of the B-line blastomeres.

**Fig. 3.**
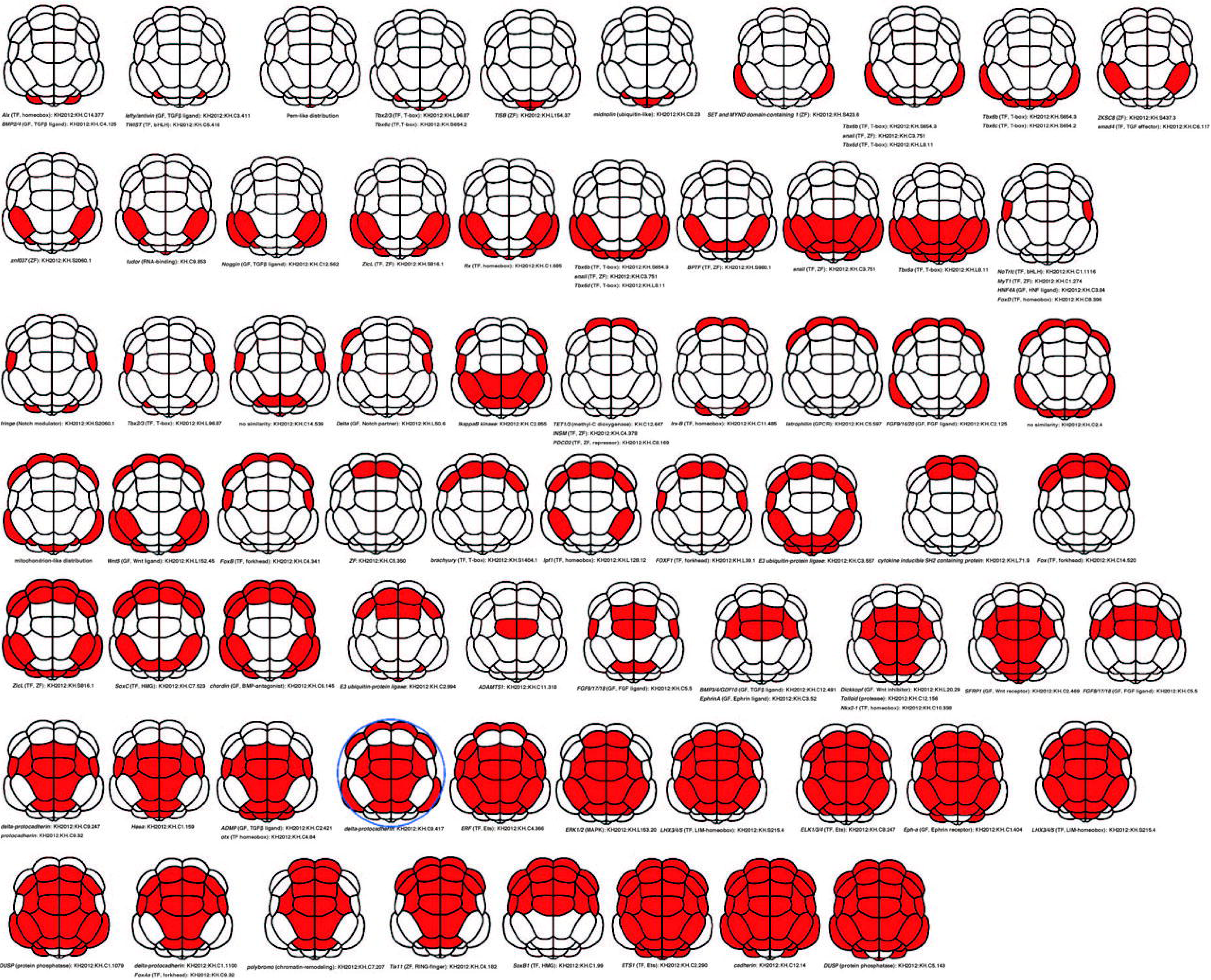
Digitized image in the Encoder program of the spatial gene expression pattern in a 64-cell *Ciona intestinalis* embryo. Sixty-eight spatial expression patterns of 89 genes in 64-cell *C. intestinalis* embryos. bHLH: basic helix-loop-helix; BMP: bone morphogenetic protein; GF: growth factor; HMG: high mobility group; FGF: fibroblast growth factor; MAPK: mitogen-activated protein kinase; RING: really interesting new gene; TF: transcriptional factor; TGFβ transforming growth factor-β and ZF: zinc finger. A blue circle indicates an embryo with blastomeres whose cell adhesions are relaxed temporally (see text).

**Fig. 4.**
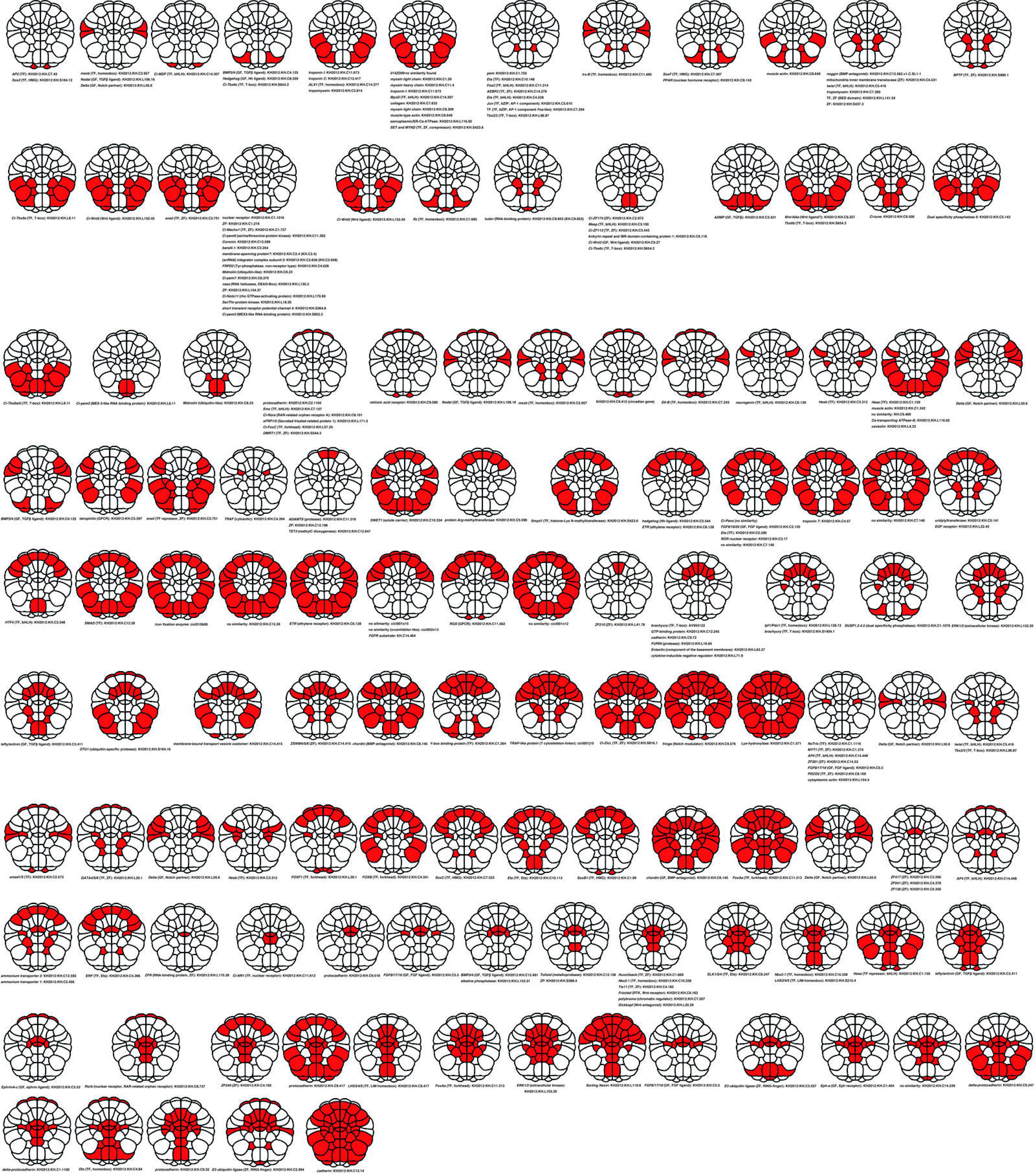
Digitized image in the Encoder program of the spatial gene expression pattern in a 110-cell *Ciona intestinalis* embryo. One hundred and twenty-one spatial expression patterns of 211 genes in 64-cell *C. intestinalis* embryos. BED: *Drosophila melanogaster* BEAF and DREF-type zinc finger; bZIP: basic leucine zipper; DEAD-box: Asp-Glu-Ala-Asp-motif-containing RNA helicase; EGF: epidermal growth factor; ETR: ethylene receptor; GPCR: G protein-coupled receptor; Hh: hedgehog; methyl-C: 5-methylcytosine; RAR: retinoic acid receptor; RTK: receptor tyrosine kinase; in addition to the abbreviations shown in Fig. 3. Red circles show different expression patterns revealed by whole-mount *in situ* hybridization experiments with the same probe.

For the analysis, I defined the expression pattern of each gene in the 16-cell embryo when the zygotic expression of most genes begins, as the provisional start of zygotic development. I also defined the expression pattern of each gene in the 110-cell embryo, when the prospective fate of most blastomeres is specified, as the provisional goal of zygotic development (Fig. 4). Between the 16-cell and 110-cell embryo, each blastomere cleaves three times.

### Four mechanisms that can establish spatial patterns of gene expression

Spatial patterns of gene expression might be established in a number of different ways during early development. It is possible to envisage that the expression patterns of some genes are determined via a two-step (rough and fine) mechanism. The segment-polarity or pair-rule genes, such as *even-skipped*, are expressed in a constant pattern of seven stripes in *D. melanogaster.* The promoter of the *even-skipped* gene, is composed of several elements; furthermore, each element is composed of up to 10 transcription-factor-binding sites (Borok et al., 2010; Fig. 5A). However, these characteristic expression patterns might be of secondary importance for development of the anterior–posterior axis in *D. melanogaster*, and thus classified as a “fine” mechanism. A pattern of seven stripes is similar to a pattern that can be found in a stationary wave, which always appears when a reflected wave is superimposed on the original wave (Aranda et al., 2008; El-Sherif et al., 2012). The seven stripes might be caused by an unknown stationary wave of gene expression (perhaps a “rough” regulator).

**Fig. 5.**
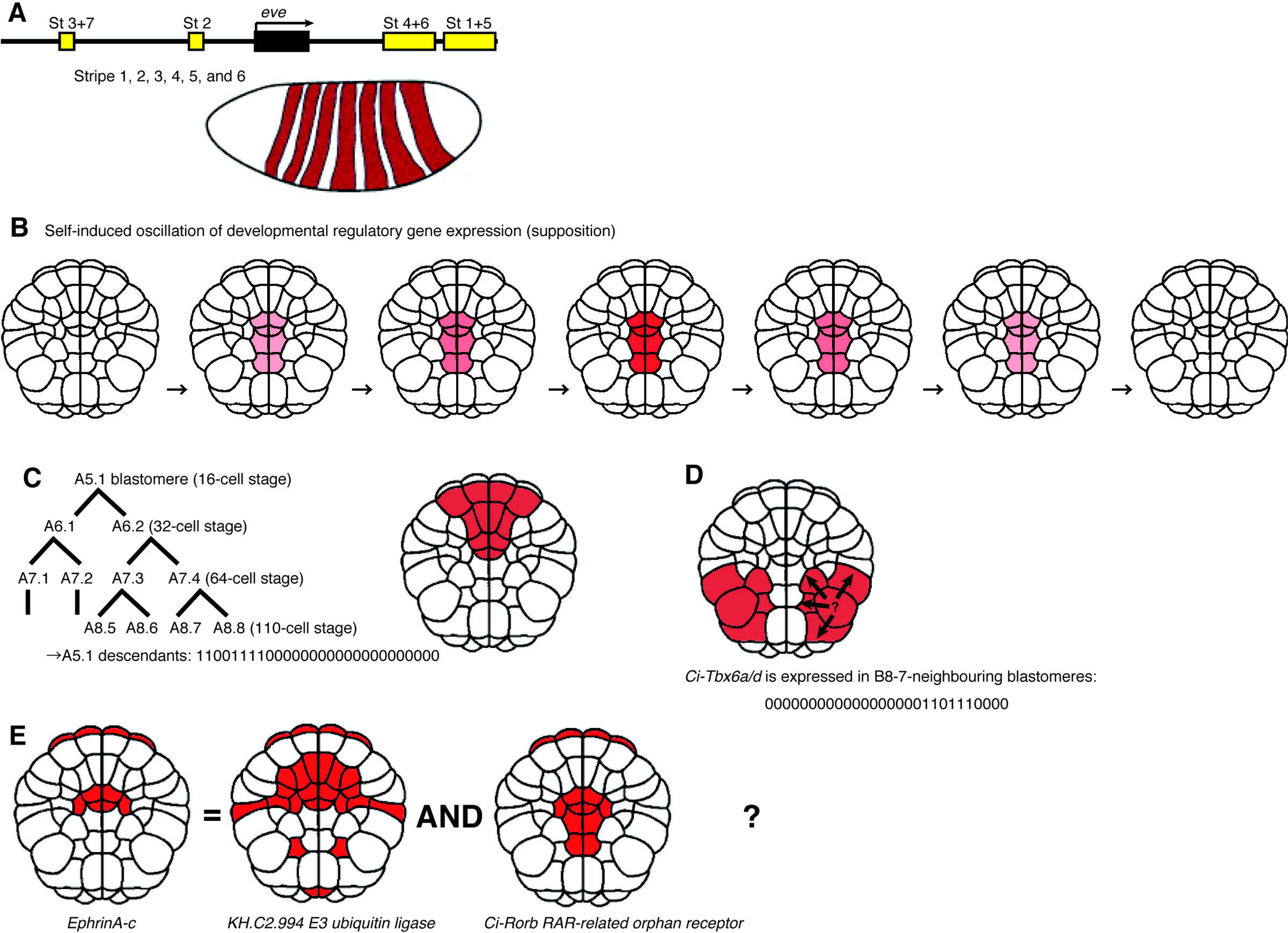
Four spatial cues for gene expression. (A) Several different regions of the promoter of the *Drosophila melanogaster even-skipped* pair-rule gene control its spatial expression pattern, giving rise to seven stripes. (B) Type 1 (no cue): expression of a temporally oscillating gene (e.g. a circadian gene). It is possible that the seven bands shown in Fig. 4A might have been caused originally by a type of stationary wave of the products of some upstream regulatory genes with temporally oscillating expression. Stationary waves arise when a wave and its reflected wave collide because the two waves have the same frequency, wavelength, and amplitude but travel in opposite directions. (C) Type 2: a maternal determinant is tethered to a certain intracellular structure in a fertilized egg (a one-cell embryo). The determinant will be inherited by the specific blastomere descendants that derive from this region of the egg. In this situation, the expression pattern is determined by the lineage of the blastomere. (D) Type 3: gene expression occurs only in blastomeres that neighbour a blastomere that is producing a certain signal molecule. In this situation, the expression pattern is determined by the geospatial (anatomical) information with regard to the blastomere. (E) The interactions among multiple upstream factors determine the expression of their downstream targets.

On the basis of the above assumption, there are several strategies by which spatial patterns of gene expression can arise (Satoh 2013; Oda-Ishii et al, 2016) and evidence of these strategies was observed in the patterns detected using the Encoder and Decoder tools.

(1) For some genes, a single probe used for whole-mount *in situ* hybridization yielded multiple spatial expression patterns, i.e. expression in a given blastomere varied between embryos. If human error is discounted and if the *in situ* hybridization images reflect the true expression at a specific time point, the different patterns might reflect multiple snapshots of appearing, disappearing or oscillating temporally dynamic gene expression (Fig. 5B). Some clock genes reportedly show oscillating expression dynamics (El-Sherif et al., 2012). If this is correct, then it would also be important to consider expression that varies temporally.

(2) A maternal factor that has been localized directly to a specific region of the fertilized egg can drive the expression of downstream target genes in the zygote. In the fertilized *C. intestinalis* egg, several maternal factors are tethered to a specific location within the egg by the centrosome-attracting body (CAB) (Hibino et al., 1998). There were no examples of genes that showed a spatial expression pattern that accurately followed the lineage information of each blastomere earlier than the 64-cell stage. This observation suggests that maternal factors do not affect such precise spatial information at the level of individual cells. Localized maternal factors in a fertilized egg might affect spatial patterns of gene expression at a broader spatial level (Fig. 5C).

(3) Induction of gene expression might depend on the geospatial arrangements of ≥2 blastomeres, for example, expression of a given gene might be activated by a signal produced by a neighbouring blastomere. For example, the A7.1 blastomere in the 110 cell-stage embryo is neighboured by A7.2, A7.5, B7.1 and another A7.1 in the opposite half of the embryo. I analysed the spatial expression patterns of each gene in relation to the geospatial characteristics of the blastomeres. However, I found little coincidence except in one case: the *Tbx6a/d* transcription factor gene is only expressed in blastomeres at the 110-cell stage that neighbour B8-7 (B8.7, B7.7, B8.5, B8.8 and B8.15; Fig. 5D).

(4) The remaining possibility is that zygotic spatial expression patterns are determined by interactions among multiple upstream regulators. It is difficult to determine how many upstream regulators exist for a certain gene. When considering the number of upstream regulators, researchers can exclude factors with a broad DNA-binding specificity from consideration. For the 211 genes under consideration, if each gene is controlled by two regulators, there are 44,521 (i.e. 211^2^) possible results in which two genes are coexpressed. If the number of regulators is more than two, includes some repressors, or the target spatial expression pattern comprises multiple results of logical operations, the human brain cannot solve this puzzle and requires the help of a computer program. Fig. 5E shows an example of a barely imaginable AND logical operation for two genes. Although this example might not represent the actual situation because it is not backed up by any *in vivo* or *in vitro* developmental biology data, it illustrates that spatial patterns of gene expression can represent the sum of such logical operations.

### Relationship between blastomeres in a 110-cell Ciona intestinalis embryo

Fig. S1 shows a dendrogram of 211 genes that is based on their spatial expression patterns. The two blastomeres b8.17/b8.19 form a protruded swelling that is known to play a central role in the development of left–right asymmetry (Nishide et al., 2012). Analysis of the images in the ANISEED database showed that KH2012:KH.C2.957 (*msxb*: *muscle segment homeobox-b*), KH2012:KH.L106.16 (*Nodal*) and KH2012:KH.L50.6 (the Notch ligand *Delta2*) are expressed in this area, which suggests the involvement of *Nodal/Notch* signaling.

I also drew a similar dendrogram to analyse the relationships among blastomeres in the 110-cell *C. intestinalis* embryo with respect to gene expression (Fig. 6). Four primordial muscle blastomeres (B8.7, B8.8, B8.15 and B8.16) showed similar patterns of gene expression and could be treated as almost identical. However, the dendrogram showed that there were some differences among these four blastomeres, for example the *chordin* gene is expressed only in B8.7 and B8.8, and not in B8.15 or B8.16.

**Fig. 6.**
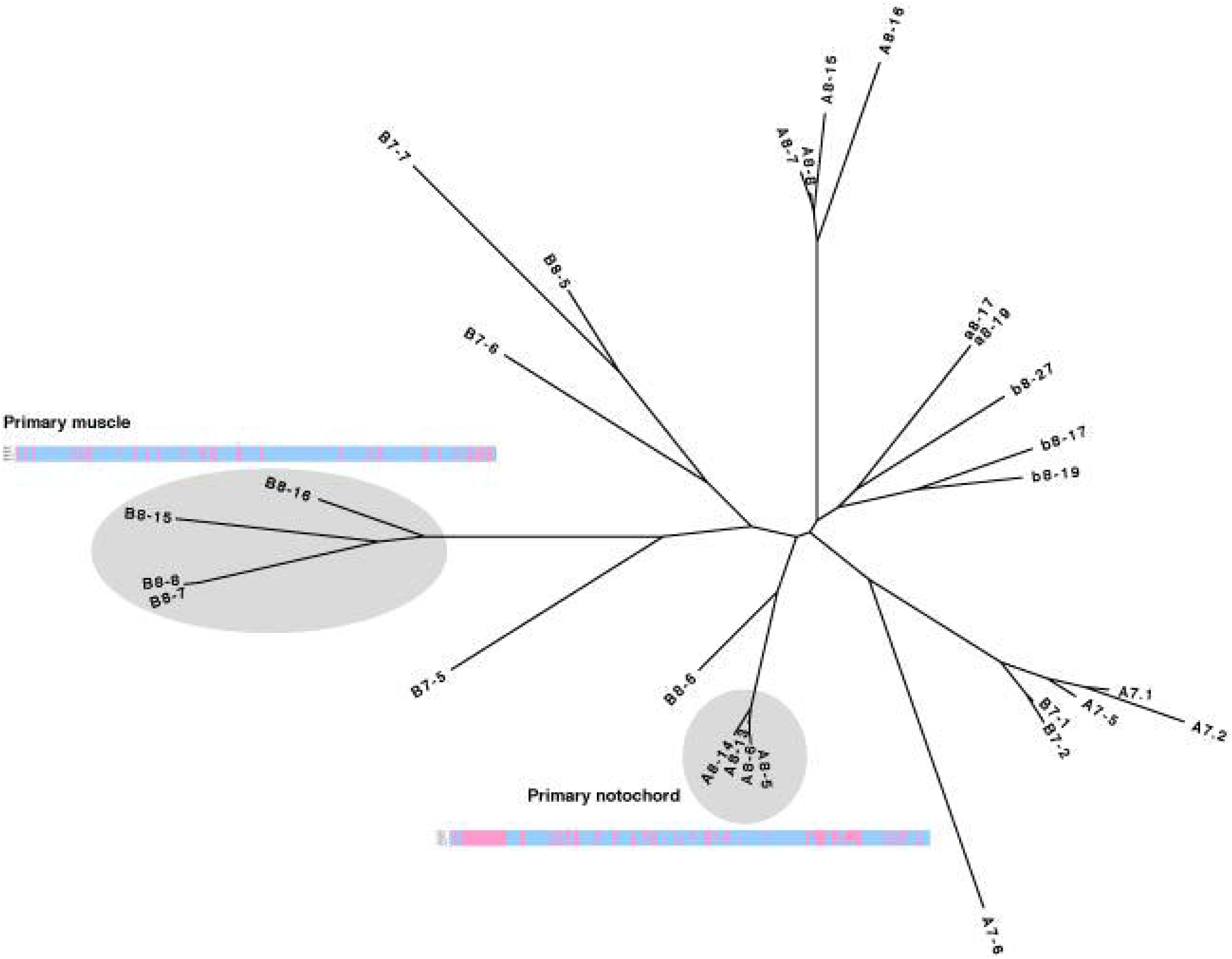
Analogous relationship of vegetal blastomeres of a 110-cell *Ciona intestinalis* embryo. A dendrogram that shows the analogous relationship of vegetal blastomeres of a 110-cell *C. intestinalis* embryo. Two diagrams show the overall similarities but subtle differences among primary muscle (upper) and primary notochord (lower) precursor blastomeres, respectively. For example, among four primordial muscle blastomeres (B8.7, B8.8, B8.15 and B8.16), only B8.7 and B8.8 blastomeres express the *chordin* gene.

### *Comparison of the data obtained* in silico *with those obtained* in vivo *and* in vitro

Detailed information is available on the regulators that bind to the promoter of *Ciona Brachyury* (*Ci-Bra*) at the cellular (blastomere) level (Ikeda and Satou, 2017). The regulators of *Ci-Bra* include four activators (*FoxAa*, *FoxD*, *ZicL* and *FGF 9/16/20*) and one repressor (*EphrinA-a*; Fig. 7A). Fig. 7B illustrates their expression patterns. The expression patterns of the *FoxAa* and *ZicL* genes differed between individual embryos (rectangles in Fig. 7B). As suggested above, these differences might represent individual captured images of temporally dynamic *FoxAa* and *FoxD* expression near the blastopore (from more central to more peripheral regions or vice versa).

**Fig. 7.**
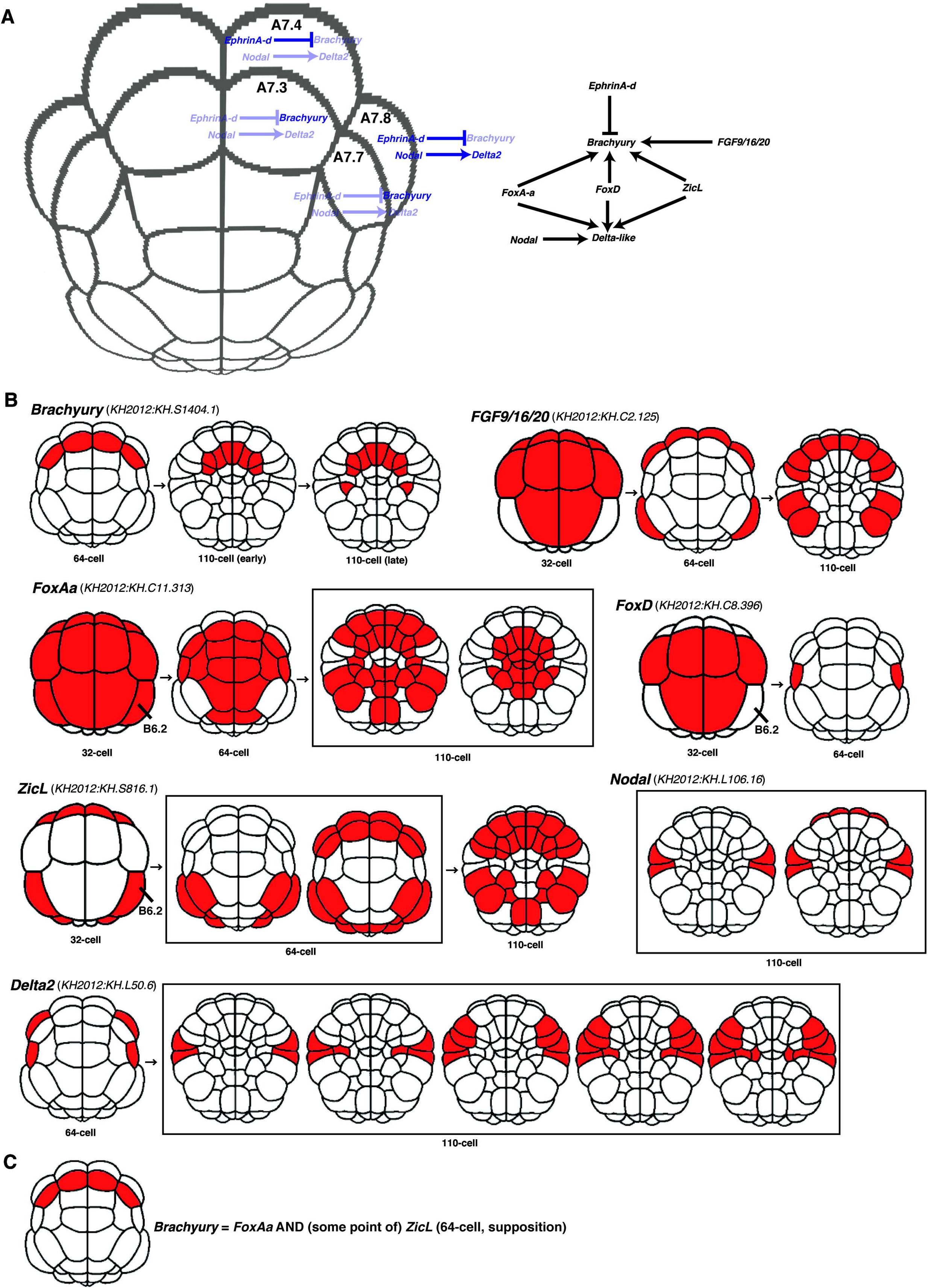
Transcriptional regulation of the *Ciona Brachyury* gene. (A) A diagram of the molecular mechanism of regulation of *Ciona Brachyury* (*Ci-Bra*) gene expression derived from the molecular developmental biology experiments of Ikeda and Satou (2017). (B) Spatial expression patterns of *Ci-Bra* and its upstream regulators. (C) *Ci-Bra* is expressed in those cells in which both *FoxAa* AND *ZicL* are expressed.

Of the four *Ci-Bra* transcriptional activators, *FoxD* was not expressed in the *Ci-Bra*-expressing blastomeres in the 64-cell embryo; however, it was expressed in their mother cells in the 32-cell embryo. Namely, *Ci-Bra* is expressed in cells in which *FoxAa* and *ZicL* are expressed in the 64-cell embryo and where *FoxD* was expressed in the mother cells in the 32-cell embryo (Fig. 7C). Although expression of *FoxAa* and *ZicL* overlapped in the B6.2 blastomere of the 32-cell embryo, *Ci-Bra* was not expressed in this blastomere, perhaps because expression of the *FoxD* regulator remained OFF (Fig. 7B).

## DISCUSSION

### *Towards elucidation of a developmental program* in silico

Although 211 genes were examined in this study, the total number of genes in *C. intestinalis* is ∼15,000. This includes 670 transcription factors, many of which belong to the zinc-finger class (Imai et al., 2004). When we include intercellular signaling-governing genes in addition to transcription factors, the number of *C. intestinalis* developmental regulatory genes is ∼1,000. Additional whole-mount *in situ* hybridization images are required to elucidate the complete *C. intestinalis* developmental program.

To decipher the developmental program, these gene expression data should perform a similar role to a latitude running in the east–west direction, i.e. they do not provide the complete picture.

As the combination of latitude and longitude specifies the position of any location on the surface of Earth, the combination of mRNA spatial distribution pattern of the gene (presence, absence or area of expression) in cellular level and biochemical activity of its product will describe the gene function sufficiently. *Ci-otx* homeobox-type transcription factor encoding gene expresses in A7.1, A7.2, A7.5, A8.11, A8.12, B7.1, B7.2, B7.5, B7.7 and B8.5 blastomere pairs of 110-cell embryo. Ci-otx gene product binds to TAATCC/T consensus DNA sequence. This combination of two lines of biological information can specify the function of *Ci-otx* gene.

Although promoter analysis studies might provide us with “longitude” information that is independent from “latitude” gene (mRNA) spatial expression pattern information, many promoter analysis studies performed to date have contained excessive mathematical information. When researchers aspire to utilize these data for “longitude” purposes, they require data on biochemical interactions between transcription factors (trans-acting factors) and promoter DNA sequences (cis-acting factors) under normal biochemical/physiological conditions with an intact nucleus (e.g. Hotta et al., 2000 and Kusakabe, 2005). Despite this, promoter analysis papers usually do not provide a complete understanding of all the factors that bind to the promoter. The case of the *Ci-Bra* gene, for which there is sufficient information, is a good exception (Fig. 7).

In this study, I used only binarized data (ON or OFF) for gene expression states. The expression level of a gene is determined by several factors, among which two are the most important: the frequency of transcriptional initiation and the lifespan of the mRNA (Jackson et al., 2000). Recently, a novel molecular mechanism, transcriptional bursting, was described; transcription occurs in discontinuous bursts and the level of gene expression depends on the frequency of the bursts. This might suggest that the molecular basis of the quantitative difference is controlled by the frequency of transcriptional initiation events, which are regulated by each qualitative state (e.g. activator A is ON, B is OFF and so on) at the promoter (Fukaya et al., 2016).

The limitations of using abnormal expression or gene knockout to examine pinpointed sites also remain to be determined. In *Halocynthia roretzi*, Miya et al. (1994) previously attempted to isolate the B4.1 blastomere of an eight-cell embryo, which should contain muscle determinants in ascidians. However, they did not isolate the *macho-1* maternal mRNA, which is the true muscle determinant as revealed in a later study (Nishida and Sawada, 2001). In *C. intestinalis*, *Hes (hairy and Enhancer of split).a* was knocked down only in the pinpointed a-line lineage, retaining the normal status of A- and B-lineages where *Hes* gene functions earlier (Ikeda and Satou, 2017). In mice, the Cre–Lox conditional gene-mutagenesis technique has been developed (Nagy, 2000), which enables site-specific recombinase technology to be used to introduce modified genes instead of the microinjection that is necessary with ascidian embryos. Such types of experimental or sophisticated molecular developmental biological techniques require the skill of an artisan and complicate additional tests by other researchers.

Dry simulations can be a substitute for wet (*in vitro* and *in vivo*) experiments to a certain extent. In particular, when there are considerable differences in gene expression among multiple blastomeres in a single developmental territory, as shown in the present study, *in vivo* or *in vitro* approaches to study the developmental program become difficult or almost impossible. It is insufficient to describe the mutant phenotype on the basis of morphological abnormalities, and the analysis of a limited number of molecular markers can improve the situation only slightly. In addition, such experiments require technical expertise. Dry simulations *in silico* can deputize for the (sometimes impossible) *in vivo* or *in vitro* experiments, although much time and effort is required to obtain the initial description of normal development. As an example, it might be possible to replace some conventional experiments, such as differential screenings or subtraction, with *in silico* experiments that, for instance, use digital differential displays of two large transcriptome data sets (Murray et al., 2007). Under such circumstances, the accuracy of each item of *in vivo* or *in vitro* data becomes more important than ever. Any uncertain data should be excluded.

If disagreement remains between the results of the analysis of the digitalized patterns of gene expression and the known molecular mechanisms of gene regulation, alternative regulatory mechanisms should be considered, such as translational regulation or post-translational mechanisms. For example, it is possible that expression of the *FoxD* activator in the 64-cell embryo is carried over from the mother cells (Fig. 7B), because notochord precursor blastomeres at the 64-cell stage do not express the gene but the mother cells at the 32-cell stage do.

### *Two morphogenetic movements during early development in* Ciona intestinalis

The morphogenesis of *C. intestinalis* is static compared with the dynamic development that is exhibited by the sea urchin or frog. In sea urchin and frog development, novel cellular interactions appear during gastrulation. Conversely, the gastrulation of *C. intestinalis* involves only the formation by the archenteron of a shallow dent (Fig. 8A).

**Fig. 8.**
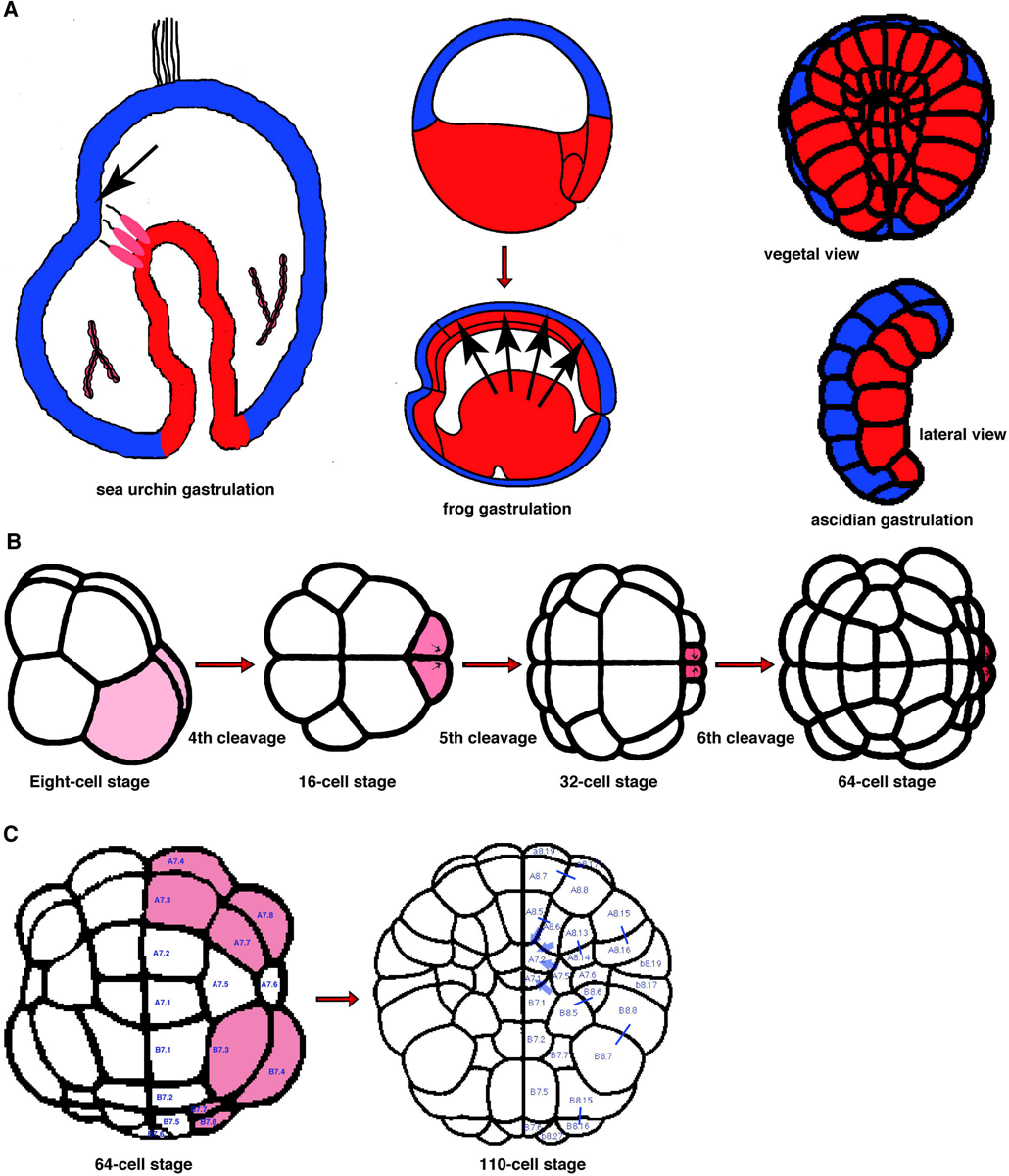
Two morphogenetic events in *Ciona intestinalis* during early embryogenesis. (A) Morphogenesis during *C. intestinalis* gastrulation (right) is relatively static compared with that in the sea urchin (left) and frog (middle). The arrows indicate newly emerged cell–cell interactions in association with the physical proximity of two cell layers. (B) The first morphogenetic movement in *C. intestinalis* development. The *C. intestinalis* embryo undergoes three successive unequal cleavages. This is caused by a tethered centrosome-attracting body (Hibino et al, 1998) that plays tug of war with the mitotic apparatus (shown in shaded blastomeres). (C) The second morphogenetic movement in *C. intestinalis* development (supposition). During *C. intestinalis* gastrulation, the synchrony of cleavages is disturbed by an unknown autonomous mechanism from the 64-cell stage; the blastopore-surrounding blastomeres (shaded) cleave before the blastopore blastomeres in the midline. This might increase the number of blastopore-surrounding blastomeres; then, these might force the midline blastomeres into the inside of the embryo.

The molecular mechanisms that underlie the morphogenetic movements that occur from the fourth to the sixth cleavages in *C. intestinalis* involve a series of unequal cleavages: the B4.1 and B5.2 micromeres cleave unequally in the fourth cleavage; the B5.2 and B6.3 micromeres cleave unequally in the fifth cleavage; and the B6.3 and B7.6 micromeres cleave unequally in the sixth cleavage (Fig. 8B). This phenomenon in the ascidian embryo is similar to the appearance of the micromeres (fourth cleavage) and small micromeres (fifth cleavage) in the sea urchin embryo in that both arise in the most posterior regions of the embryos. In ascidians, these unequal cleavages are caused by an intracellular structure, the CAB, which contains various maternal factors and is tethered to the most posterior pole of the embryo (Hibino et al., 1998).

In addition, the synchronicity of cleavages in the ascidian embryo is disturbed after the 32-cell stage. The 110-cell embryo contains both the seventh- and eighth-generation blastomeres. Blastomeres that surround the blastopore seem to cleave earlier than those near the blastopore center. This contributes to the aforementioned disturbance of synchronicity. The blastomeres that cleaved earlier then exert force on the remaining ones. This second morphological movement might represent a cellular mechanism of static ascidian gastrulation.

If one scrutinizes the gene expression patterns of blastomeres in a 64-cell embryo, it is evident that the blastomeres at the blastopore center express the *ZicL* and *noggin* genes but lack expression of *delta-protocadherin* (KH2012:KH.C9.417, blue circle in Fig. 3). It is thought that cleaving blastomeres relax their cell-adhesion activity by turning off expression of this cadherin. Cells generally loosen adhesive cell contacts and relax temporarily just before cell division, as is seen in cancer cells. In ascidian early embryogenesis, it is possible that this change in expression is implemented at the level of transcriptional control rather than at the translational or post-translational level.

## MATERIAL AND METHODS

### Ethical Statement

The author did not carry out any wet molecular biological studies in this research.

### Digitization of spatial expression patterns of genes published in the ANISEED database

I have developed two computer programs (Encoder and Decoder) to digitalize microscope images of whole-mount *in situ* hybridization analyses (Fig. 2; Harada, 2016). For each gene analysed, a score of 1 or 0 was assigned to each blastomere depending on whether gene expression was ON or OFF. Both the Encoder and Decoder programs are freely downloadable from the figshare repository (https://figshare.com).

The three Encoder tools show vegetal images of the *C. intestinalis* embryo at the 32-, 64- and 110-cell stages respectively, when most of the whole-mount *in situ* hybridization analyses were carried out. Each blastomere has a “radio” button assigned to it. The radio button is activated by the researcher to indicate that the blastomere expresses this gene at the relevant developmental stage. The results form a character string of binary digits for each embryo examined. The Decoder shows a stack of layers of multiple blastomere images with a transparent background. It can provide a representation of expression throughout an entire vegetal image to visualize the character string.

Two hundred and seventeen microscope images of embryos in the ANISEED ascidian development browser (Tassy et al., 2010; Brozovic et al., 2016), which were annotated with the names of the genes that were expressed in each blastomere, were converted into character strings using the Encoder tool (Fig. **2A**). Some whole-mount *in situ* hybridization DIG probes gave multiple images, and consequently the expression patterns of 211 genes were digitalized. AND, OR and NOT bit calculations were performed on the data obtained (data not shown), and they were sorted into categories on the basis of information on blastomere lineage (e.g. the A5.1 blastomere at the 16-cell stage gives rise to six [12] blastomeres at the 110-cell stage) and geospatial information (e.g. the B8.7 blastomere at the 110-cell stage neighbours five [10] blastomeres: B7.7, B8.5, B8.7 itself, B8.8 and B8.15 blastomeres at the 110-cell stage) using Excel (Microsoft, Redmond, WA, USA).

### Construction of phylogenetic trees of genes and blastomeres

To construct phylogenetic trees of the expressed genes, I constructed a guide tree (*.dnd file), which consisted of a dendrogram made temporarily using the ClustalW multiple sequence alignment software (http://www.genome.jp/tools/clustalw/).

In the character strings from the Encoder program, the 1 and 0 correspond to the gene expression states ON and OFF, respectively. The 1 and 0 codes were substituted with “G” and “A”, respectively, to enable ClustalW to recognize the data. The dendrogram was drawn using the FigTree software (http://tree.bio.ed.ac.uk/software/figtree/).

In order to construct a blastomere phylogenetic tree, the expression character strings were decomposed into pieces of one character and transposed the columns and rows in the Excel worksheet.

## Acknowledgements

I thank ascidian developmental biologists worldwide who have performed the whole-mount *in situ* hybridization analyses, for example, Profs. Shigeki Fujiwara (Kochi University), Takahito Nishikata (Konan University), Michio Ogasawara (Chiba University), Takehiro Kusakabe (Konan University), and Eiichi Shoguchi (OIST), and the director of this project Prof. Nori Satoh (OIST). I also express gratitude to Prof. Patrick Lemaire (CRBM) and his colleagues for the public ANISEED database, Yutaka Satou and his colleagues for the public ghost database, which provided most of the original data for ANISEED and Hiroki Nishida (Osaka University, http://www.bio.sci.osaka-u.ac.jp/bio_web/lab_page/nishida/maboya-e.html) for digital images on ascidian development. I dedicate this paper to the memory of Prof. Eric H. Davidson (Caltech) with the deepest thanks.

### Funding

#### Competing interests

The author declares no competing interests.

### Author contributions

#### List of abbreviations and symbols used

ANISEED: Ascidian Network for *In Situ* Expression and Embryological Data
CAB: centrosome-attracting body FGF: fibroblast growth factors Fox: forkhead-box
GRN: gene regulatory network Hes: hairy and Enhancer of split
Sox: Sry-related high mobility group box Zic: zinc-finger

## SUPPLEMENTARY FIGURE

**Supplementary Figure 1. The analogous relationships among spatial gene-expression patterns in a 110-cell *Ciona intestinalis* embryo.**

